# Age differences in brain functional connectivity underlying proactive interference in working memory

**DOI:** 10.1101/2024.06.18.599495

**Authors:** P Andersson, M.G.S. Schrooten, J Persson

**Affiliations:** Center for Life-span Developmental Research (LEADER), School of Behavioral, Social and Legal sciences, Örebro University, Sweden; Aging Research Center (ARC), Karolinska Institute and Stockholm University, Sweden; Center for Health and Medical Psychology (CHAMP), School of Behavioral, Social and Legal sciences, Örebro University, Sweden

## Abstract

Aging is typically accompanied by a decline in working memory (WM) capacity, even in the absence of pathology. Proficient WM requires cognitive control processes which can retain goal-relevant information for easy retrieval and resolve interference from irrelevant information. Aging has been associated with a reduced ability to resolve proactive interference (PI) in WM, leading to impaired retrieval of goal-relevant information. It remains unclear how age-related differences in the ability to resolve PI in WM are related to patterns of resting-state functional connectivity (rsFC) in the brain. Here we investigated the association between PI in WM and rsFC cross-sectionally (n = 237) and 5-year longitudinally (n = 134) across the adult life span by employing both seed-based and data-driven approaches. Results revealed that the ability to resolve PI was associated with differential patterns of rsFC in younger/middle-aged adults (25-60y) and older adults (65-80y) in two clusters centered in the vermis and caudate. Specifically, more PI was associated with stronger inferior frontal gyrus – vermis connectivity and weaker inferior frontal gyrus – caudate connectivity in older adults, while younger/middle-aged adults showed associations in the opposite directions with the identified clusters. Whole brain multivariate pattern analyses showed age-differential patterns of rsFC indicative of age-related structural decline and age-related compensation. The current results show that rsFC is associated with the ability to control PI in WM and that these associations are modulated by age.

## 1. INTRODUCTION

Getting older is commonly accompanied by increasing memory difficulties (Koen & Yonelinas, 2014; Nyberg et al., 2020), even in the absence of any memory pathology such as dementia. An efficient memory system entails the ability to control the flow of information that we are almost constantly processing, to determine what information should be retained. Declining control processes might result in cognitive overload and impede goal-directed behavior that relies on the ability to selectively engage relevant information. Crucially, irrelevant information might interfere with goal-relevant information. *Proactive interference* (PI) occurs when old, outdated information interferes with memory for new information, leading to forgetting (Hasher & Zacks, 1988; Jonides & Nee, 2006).

While PI has traditionally been studied primarily in the context of long-term episodic memory (Keppel, 1968; Roediger & McDermott, 1995; Kliegl & Bäuml, 2021), evidence supports its presence also in working memory (WM; Bunting, 2006; Emery, Hale, & Myerson, 2008). PI and the inability to appropriately control PI have been considered a major source of forgetting, not only in long-term memory (Kliegl & Bäuml, 2021) but also in WM (Oberauer & Lewandowsky, 2008). Resolving PI in WM has been consistently localized to a network of brain regions consisting of the left inferior frontal gyrus (IFG), the anterior cingulate cortex, and the striatum/insula (Badre & Wagner, 2005; Burgess & Braver, 2010; Jonides & Nee, 2006; Marklund & Persson, 2012; Nelson et al., 2009; Persson, Larsson, & Reuter-Lorenz, 2013; Samrani, Bäckman, & Persson, 2019).

An age-related decrease in the ability to control PI has been found in both long-term memory (Healey et al., 2013; Wahlheim, 2014) and WM (Loosli et al., 2016; Samrani & Persson, 2021). Furthermore, age-related differences in cognition, such as episodic memory, are largely explained by PI in WM, over and above the effects of processing speed (Samrani & Persson, 2021).

Increasing age is characterized by changes in both the structure and function of the brain. Specifically, aging is associated with a decrease in gray matter volume (Fjell et al., 2014) as well as reduced white-matter integrity (Sexton et al., 2014). These structural changes have been postulated to underly age-related functional changes in the brain (Greenwood, 2007; Park & Reuter-Lorenz, 2009). Functionally, increasing age is associated with neural dedifferentiation, which is identified by less selective neural processing and might explain reduced neural efficiency with age (Koen, Joshua, & Rugg, 2019; for review: Goh, 2011). Moreover, older adults tend to have decreased activity in posterior regions and increased activity in frontal regions, compared to younger/middle-aged adults (Davis et al., 2008). It has been suggested that this pattern may be associated with less efficient functional communication, reflected by reduced resting state functional connectivity (rsFC) between networks (Goh, 2011). Older adults also display less functional connectivity with posterior regions and more connectivity with frontal regions compared to younger/middle-aged adults (Zhang, Lee, & Qiu, 2017). An age-related increase in frontal activity together with the recruitment of additional regions or bilateral activation has been associated with maintained cognitive performance including WM (Goh, 2011). This pattern is consistent with the Compensation Related Utilization of Neural Circuits (CRUNCH) model (Reuter-Lorenz & Cappell, 2008), which proposes that older adults compensate for deficient neural resources through additional functional recruitment. Moreover, weaker rsFC in older adults has been associated with poorer executive functioning, processing speed (Damoiseaux et al., 2008), and associative memory (Wang et al., 2010).

The structural and functional neurological changes that occur with aging may underly age-related decline in the ability to control PI.. Specifically, more PI in older adults has been associated with smaller gray-matter volume in the IFG (Samrani et al., 2019) and the hippocampus (HC; Andersson et al., 2023), lower white-matter integrity (Andersson et al., 2022), altered task-related BOLD activation (Loosli et al., 2016), and altered functional connectivity (Oren et al., 2017). Oren et al. (2017) found that whereas greater connectivity between the (left) IFG and the hippocampi predicted PI in all participants, PI was more related to inter-subject connectivity in the posterior cingulate cortex (PCC) in older adults. Importantly though, this study did not employ a WM task to investigate PI but studied PI as context-effects of successive movies on a linguistic task.

A few other studies have investigated functional connectivity in relation to control of PI, though without considering age-related differences. For example, Samrani and Persson (2022) found that a longer interval between target and lure items in an n-back task, which results in lower PI, was associated with increased connectivity between HC and IFG, insula, ACC, thalamus, putamen, and superior temporal regions. Using the recent probes and directed-forgetting tasks, Nee et al. (2007) found significant connectivity between IFG with the premotor cortex, medial temporal cortex, anterior cingulate cortex, inferior temporal pole, PCC, and caudate and between the anterior prefrontal cortex with the anterior cingulate cortex in response to PI. Collectively, these results highlight the IFG, a region suggested to support PI resolution (Badre & Wagner, 2005; Burgess & Braver, 2010; Jonides & Nee, 2006; Marklund & Persson, 2012; Nee, Wager, & Jonides, 2007; Nelson et al., 2009; Persson, Larsson, & Reuter-Lorenz, 2013; Samrani, Bäckman, & Persson, 2019; Samrani & Persson, 2024; Öztekin et al., 2008) as an important node in the functional connectivity network associated with controlling PI. Despite age-related differences in rsFC, there are to our knowledge no previous studies investigating rsFC patterns associated with PI in WM in the context of aging. Such investigations can provide novel insights into the intrinsic organization of brain networks underlying the ability to control PI that may not be evident in task-based or structural imaging and could thus improve our understanding of network alterations underlying age-related decline in WM and more specifically PI control.

In this study, we investigated rsFC patterns associated with PI in a large (Nbaseline = 237) population-based, longitudinal (5 years) dataset using an interference version of the n-back WM task. First, we investigated the rsFC between an established node of the brain network associated with controlling PI in WM (namely IFG) and the rest of the brain, and how this connectivity is related to PI, across the whole sample as well as dependent on age. Secondly, using data-driven multivariate pattern analysis we explored age-related differences in whole-brain connectivity patterns associated with PI, separately for individuals scoring low and high on PI. These approaches can provide unique and novel insights into the network dynamics associated with PI, as well as age-related ability to appropriately control PI. Based on previous findings, we expected to find (1) weaker IFG rsFC in relation to increasing age; (2) alterations in whole brain rsFC patterns in relation to the ability to control PI in WM; and (3) that weaker IFG connectivity with other regions within the PI network would be associated with a lower ability to control PI in older adults.

## 2. MATERIALS AND METHODS

### 2.1 Participants

Participants were recruited from *The Betula prospective cohort study: Memory, health, and aging* (Nilsson et al., 1997; Nyberg et al., 2020), a deeply phenotyped longitudinal cohort. Participants were included from samples for which MRI measures were collected in 2008-2010 (timepoint 5, baseline) and 2013-2014 (timepoint 6, follow-up). The Betula study was approved by the Regional Ethical Review Board in Umeå (dnr: 2008-08-132 and dnr: 2013-92-31M), and written consent was obtained from every participant.

Individuals with clinical dementia and other neurological disorders at baseline or follow-up were excluded from the current analyses. Dementia status was assessed at baseline and reassessed every 5 years using a three-step procedure. First, an overall evaluation was performed by an examining physician according to the Diagnostic and Statistical Manual of Mental Disorders, 4th edition (DSM–IV; American Psychiatric Association, 1994). Second, using a composite measure based on scores from several cognitive tests (episodic memory, WM, processing speed, semantic memory, and fluid intelligence), each participant was compared to the mean cognitive score for their age cohort. If an individual scored more than two standard deviations below the mean of their age cohort, they would be flagged for further assessment of dementia by a clinical psychiatrist. Third, all participants scored at or above the cut-off score for dementia of 24 using MMSE (Mini Mental State Examination, Folstein et al., 1975). To retain the diversity of the sample, exclusions were not made for: diabetes, hypertension, mild depressive symptoms, and other moderately severe medical conditions, which are common in older participants.

Participants with extremely low performance on the n-back task (proportion hits minus proportion false alarms < .1), indicating a very low adherence to task instructions, were excluded (17 participants). Six additional participants who did not pass these performance-based criteria at baseline but passed at the second time-point were included at follow-up testing. The final total sample consisted of 237 participants for cross-sectional analyses at baseline (25-80 years, M = 58.9, SD = 13.5, M education = 13 years (SD = 4 years); Supplementary Figure 1), and 134 participants for longitudinal analyses at follow-up. For a drop-out analysis, see Noroozian et al. (2023). Demographic information and cognitive performance can be found in Table 1.

**Table 1.**
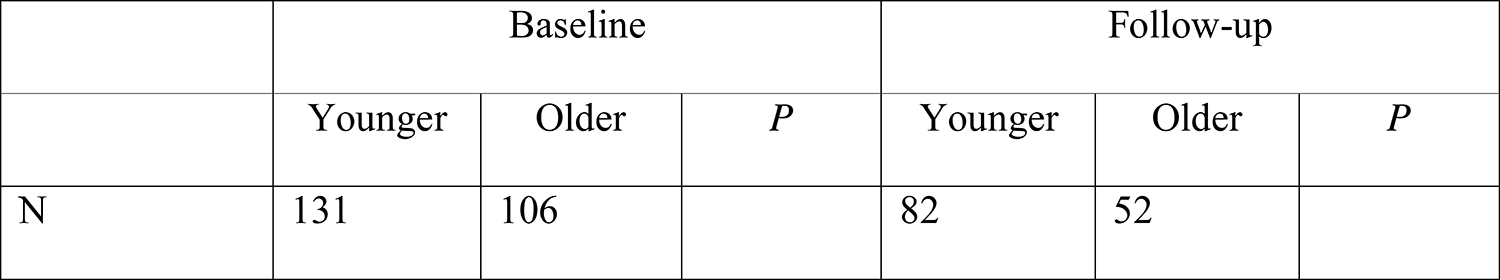
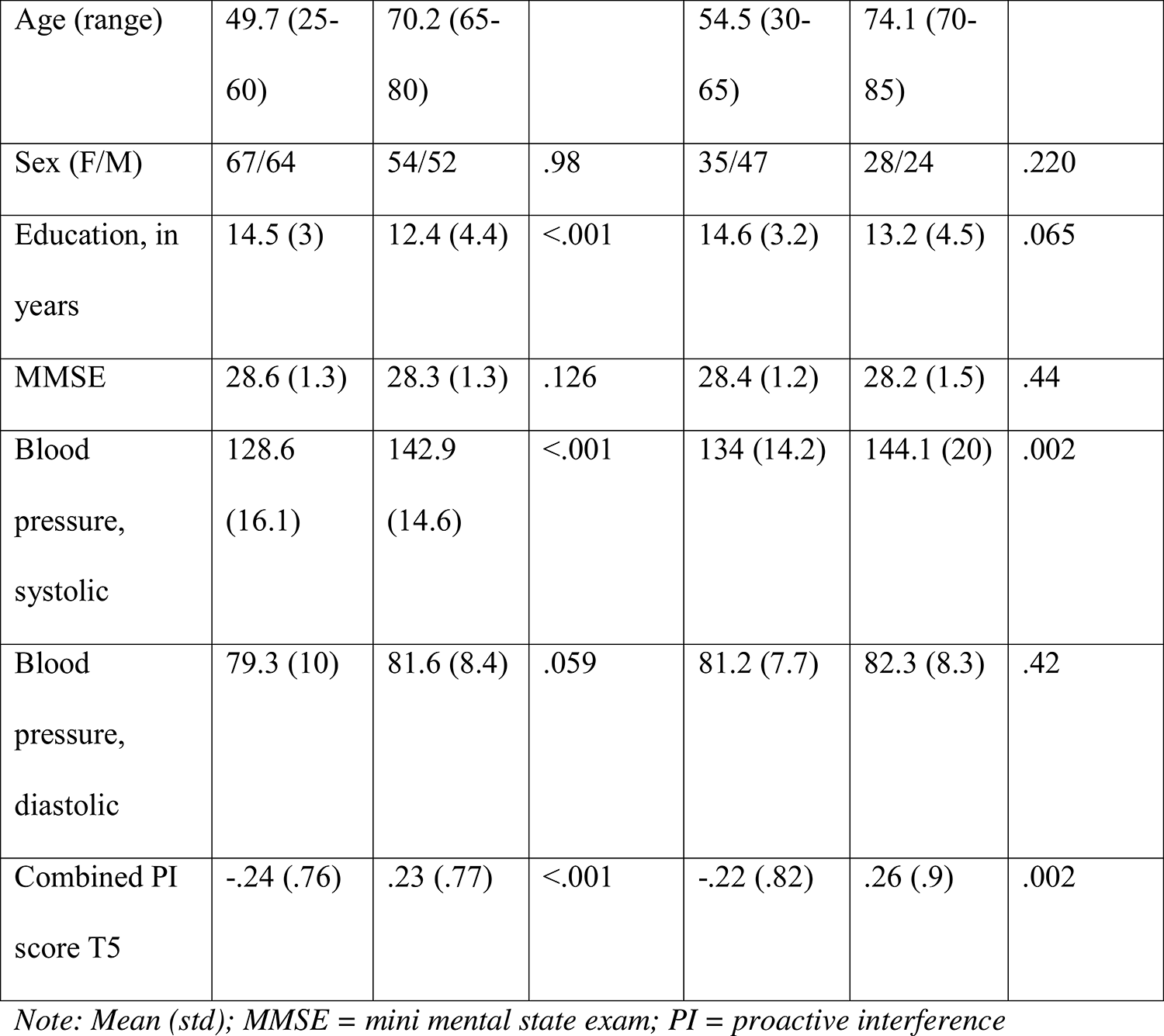
Sample demographical and cognitive information.

### 2.2. Cognitive measures

PI in WM was measured using a modified version of the verbal 2-back task which was designed to induce proactive interference (Gray, Chabris, & Bravor, 2003; Marklund & Persson, 2012; Nee et al., 2007). Participants were presented with a series of Swedish nouns, one after the other, and instructed to indicate for every noun whether it matches the one presented two trials prior. On *target trials*, the current stimulus matches the one presented two trials earlier (requiring a ‘Yes’ response); on *new trials*, the current stimulus has not been presented in previous trials, and so is non-familiar (requiring a ‘No’ response); and on *lure trials*, the current stimulus matches a stimulus presented 3- or 4 trials prior, and so is familiar (requiring a ‘No’ response). The task consisted of 40 trials (9 target trials, 21 new trials, 8 3-back lure trials, and 2 4-back lure trials). All stimuli and trial conditions were presented in the same fixed random order to all participants. Stimuli were presented for 2500 ms (ITI = 2000 ms). Participants were instructed to respond as quickly and accurately as possible by pressing the “m” key for ‘yes’ and the “x” key for ‘no’, on a standard Swedish qwerty keyboard, using their right and left index finger, respectively.

RT calculations were based on correct responses (in milliseconds). To reduce the influence of extreme values, median RTs were used. Accuracy was estimated as percentage of correct responses out of total number of trials, excluding omissions.

PI scores were calculated by combining the relative proportional difference in RT and accuracy between new trials (requiring ‘No’ responses to non-familiar stimuli) and lure trials (requiring ‘No’ responses to familiar stimuli). First, a relative difference score ((*lure trials/new trials*−1) ×100) was calculated for accuracy and RT separately. Using a relative difference score should provide a more salient and individual measure of executive control, taking into consideration the individual, age-related differences in the variable. Relative difference scores based on RT and accuracy were positively correlated at baseline, *r*(237) = .299, *p* = < .001, and follow up, *r*(132) = .330, *p* = < .001. Second, the two scores were normalized with z-score transformation. Finally, the scores were combined in a PI score by calculating the average of the two difference scores. Higher PI scores indicate a lower ability to control PI.

### 2.3 Procedure

Scanning was performed at the MR-center (Umeå Center for Functional Brain Imaging; UFBI) at the University hospital in Umeå, and the cognitive (2 hours) and medical (2 hours) test sessions were performed on two separate days at the Psychology department at Umeå University. The cognitive test session was performed before the scanning session for all participants. Informed consent was obtained before the cognitive test session from all participants.

### 2.4 MRI data acquisition

MRI data were collected using a 3T GE scanner (32-channel head coil). T1-weighted images were acquired with a 3D fast spoiled gradient echo sequence (180 slices; thickness = 1 mm; TR = 8.2 ms; TE = 3.2 ms; 12° flip angle; FOV = 25 x 25 cm). Resting state data were acquired with a gradient echoplanar imaging sequence (37 transaxial slices; thickness = 3.4 mm; gap = .5 mm; TR = 2 s; TE = 30 ms; 80° flip angle; FOV = 25 x 25 cm). The first 10 scans were excluded from the experimental image acquisition as dummy scans. The same scanner was used for baseline and follow-up.

### 2.5 Scanner stability

Scanner software changes were implemented from baseline to follow-up. To check whether these changes affected image quality, a quality assure program (based on Friedman & Glover, 2006) was run weekly, confirming scanner stability.

### 2.6 Statistical analyses

Based on previous demonstrations of age-differential relationships between brain function and structure in younger and older adults (Burzynska et al., 2012; Koen & Rugg, 2019; Rieckmann et al., 2018; Van Petten, 2004), whole-sample analyses were complemented with age-stratified analyses. Participants were divided into two age groups based on their age at baseline with 65 years as the cut-off age: one *younger/middle-aged group* (N = 131, 25-60 years) and one *older group* (N = 106, 65-80 years).

Based on previous demonstrations of differential relationships between brain activity and cognitive performance in older adults with preserved cognitive performance and those exhibiting cognitive decline (Damoiseaux et al., 2008; Goh et al., 2011; Wang et al., 2010), PI groups were constructed based on a median split of the PI scores (median = −.044) across age groups. The split resulted in a *low PI group* consisting of 119 individuals (39 older, 80 younger/middle-aged) displaying relatively higher performance on the n-back task indicated by less PI, and a *high PI group* consisting of 118 individuals (67 older, 51 younger/middle-aged) displaying relatively lower performance on the n-back task indicated by more PI (Supplementary Table 1). The PI groups were used in the multivariate pattern analyses (see section 2.7.2.) to investigate age-group related differences in rsFC patterns within the low and high PI groups.

Whole-sample and age-stratified partial correlation analyses were conducted to investigate the relationship between age and PI at baseline and between age and change in PI from baseline to follow-up, controlling for sex (F/M) and education level (in years). Change scores were calculated by dividing follow-up PI scores by baseline PI scores. A 2 (age group) × 2 (PI group) ANOVA was performed to evaluate the main effects of age group and PI group and the age group × PI group interaction effect on PI scores at baseline. Post-hoc t-tests were performed to compare PI scores at baseline between age-groups, within as well as between PI groups.

### 2.7 MRI preprocessing and analysis

Data were preprocessed and analyzed using CONN (Whitfeld-Gabrieli & Nieto-Castanon, 2012; RRID:SCR_009550) release 22.a (Nieto-Castanon & Whitfeld-Gabrieli, 2022) and SPM (Penny et al., 2011) (RRID:SCR_007037) release 12.7771. All reported coordinates are in MNI (Montreal Neurological Institute) space.

Functional and anatomical data were preprocessed using CONN’s default preprocessing pipeline (Nieto-Castanon, 2020), including realignment with correction of susceptibility distortion interactions, slice timing correction, outlier detection, direct segmentation and MNI-space normalization, and smoothing. Functional data were realigned using SPM realign & unwarp procedure (Andersson et al., 2001), where all scans were co-registered to a reference image (first scan of the first session) using a least squares approach and a 6 parameter (rigid body) transformation (Friston et al., 1995), and resampled using b-spline interpolation to correct for motion and magnetic susceptibility interactions. Temporal misalignment between different slices of the functional data (acquired in interleaved bottom-up order) was corrected following SPM slice-timing correction (STC) procedure (Henson et al., 1999; Sladky et al., 2011), using sinc temporal interpolation to resample each slice BOLD time series to a common mid-acquisition time. Potential outlier scans were identified using ART (Whitfeld-Gabrieli et al., 2011) as acquisitions with framewise displacement above 0.9 mm or global BOLD signal changes above 5 standard deviations (Nieto-Castanon, submitted; Power et al., 2014). A reference BOLD image was computed for each subject by averaging all scans excluding outliers. Functional and anatomical data were normalized into standard MNI space, segmented into grey matter, white matter, and CSF tissue classes, and resampled to 2 mm isotropic voxels following a direct normalization procedure (Calhoun et al., 2017; Nieto-Castanon, submitted) using SPM unified segmentation and normalization algorithm (Ashburner & Friston, 2005; Ashburner, 2007) with the default IXI-549 tissue probability map template. Last, functional data were smoothed using spatial convolution with a Gaussian kernel of 8 mm full width half maximum (FWHM).

Functional data were denoised using a standard denoising pipeline (Nieto-Castanon, 2020) including the regression of potential confounding effects characterized by white matter timeseries (5 CompCor noise components), CSF timeseries (5 CompCor noise components), motion parameters and their first order derivatives (12 factors; Friston et al., 1996), outlier scans (below 84 factors; Power et al., 2014), and linear trends (2 factors) within each functional run, followed by bandpass frequency filtering of the BOLD timeseries (Hallquist, Hwang & Luna, 2013) between 0.008 Hz and 0.09 Hz. CompCor (Behzadi et al., 2007; Chai et al., 2012) noise components within white matter and CSF were estimated by computing the average BOLD signal as well as the largest principal components orthogonal to the BOLD average, motion parameters, and outlier scans within each subject’s eroded segmentation masks. From the number of noise terms included in this denoising strategy, the effective degrees of freedom of the BOLD signal after denoising were estimated to range from 20.3 to 95.8 (average 77.1) across all participants (Nieto-Castanon, submitted).

#### 2.7.1 Seed-based analyses

Seed-based connectivity (SBC) maps were estimated characterizing the spatial pattern of functional connectivity with the IFG as seed area. Functional connectivity strength was represented by Fisher-transformed bivariate correlation coefficients from a weighted general linear model (weighted-GLM; Nieto-Castanon, 2020b), estimated separately for the seed area and each target voxel, modeling the association between their BOLD signal timeseries.

The IFG was selected as a region of interest (ROI) in the seed-based analyses based on previous evidence suggesting this region may support cognitive control processes involved in regulating PI (Badre & Wagner, 2005; Burgess & Braver, 2010; Jonides & Nee, 2006; Marklund & Persson, 2012; Nee, Wager, & Jonides, 2007; Nelson et al., 2009; Persson, Larsson, & Reuter-Lorenz, 2013; Samrani, Bäckman, & Persson, 2019; Samrani & Persson, 2024; Öztekin et al., 2008). The IFG was anatomically defined by including AAL subregions pars opercularis, pars triangularis, and pars orbitalis. To reduce the number of comparisons, and because there were no hypotheses regarding laterality or sub-regional specificity, a binary mask of the IFG was created using the WFU Pickatlas toolbox (AAL atlas; RRID:SCR_007378), combining the left and right hemisphere.

Seed-based connectivity correlational analyses were conducted to investigate the main effect of PI on IFG rsFC at baseline with the between-subject contrast (all_participants, PI_baseline [0 1]). Age-group differences in PI-related IFG rsFC at baseline were examined with a between-subject contrast of (older, younger/middle-aged, older × PI_baseline > younger/middle-aged × PI_baseline [0 0 1 −1]). Change in IFG rsFC over five years (baseline– follow-up) was examined with a within-subject contrast of (rsFC baseline < rsFC follow-up [-1 1]). Age-group differences in the relationship between change in PI and change in IFG rsFC over five years (from baseline to follow-up) were examined with a between-subject contrast of (older, younger/middle-aged, older × PI_change > younger/middle-aged × PI_change [0 0 1 −1]. A voxel-wise threshold of *p* = .05 and a cluster-level threshold (FDR-corrected) of *p* = .05 were used to identify significant clusters. Participant-specific mean cluster connectivity values from clusters identified in the seed-based analyses were extracted and imported to SPSS (v.26; IBM, Armonk, NY, USA) for further visualization and analysis.

#### 2.7.2 Multivariate pattern analyses (MVPA)

Functional connectivity multivariate pattern analyses (fc-MVPA; Nieto-Castanon, 2022) were performed to estimate the first 12 eigenpatterns characterizing the principal axes of heterogeneity in functional connectivity across participants and time points. From these eigenpatterns, 12 associated eigenpattern-score images were derived for each individual subject and condition characterizing their brain-wide functional connectome state.

Eigenpatterns and eigenpattern-scores were computed separately for each individual seed voxel as the left- and right-singular vectors, respectively, from a singular value decomposition (group-level SVD) of the matrix of functional connectivity values between this seed voxel and the rest of the brain (a matrix with one row per target voxel, and one column per subject and condition). Individual functional connectivity values were computed from the matrices of bivariate correlation coefficients between the BOLD timeseries from each pair of voxels, estimated using a singular value decomposition of the z-score normalized BOLD signal (subject-level SVD) with 64 components separately for each subject and condition (Whitfeld-Gabrieli & Nieto-Castanon, 2012).

Group-level analyses were performed using a General Linear Model (GLM; Nieto-Castanon, 2020c). For each individual voxel a separate GLM was estimated, with first-level connectivity measures at this voxel as dependent variables (one independent sample per subject and one measurement per time point), and PI group (2: Low PI, High PI) × Age group (2: Older, Younger/middle-aged) as independent variables. Voxel-level hypotheses were evaluated using multivariate parametric statistics with random-effects across participants and sample covariance estimation across multiple measurements. Inferences were performed at the level of individual clusters. Cluster-level inferences were based on parametric statistics from Gaussian Random Field theory (Worsley et al., 1996; Nieto-Castanon, 2020d). Results were thresholded using a combination of a cluster-forming p < 0.001 voxel-level threshold, and a familywise corrected p-FDR < 0.001 cluster-size threshold (Chumbley et al., 2010).

We used MVPA with 12 eigenpatterns to identify differences in brain-wide functional connectivity pattern using the between-participants contrast of the younger/middle-aged vs. older group (−1 1) within the Low PI and High PI group, separately. The eigenpattern maps were analyzed simultaneously using an omnibus/*F*-test. Thus, two MVPA models were conducted in total. Twelve eigenpatterns corresponds to an approximate 1:10 eigenpattern-to-participant ratio, which is considered a relatively conservative approach (Nieto-Castanon, 2022). A voxel-wise threshold of *p* = .001 and a cluster-level threshold (FDR-corrected) of *p* = .001 were used to identify significant clusters.

##### 2.7.2.1 Post-hoc analyses

As MVPA is an omnibus test and, therefore, does not inform on the nature of identified differences in connectivity patterns, we conducted post-hoc seed-based analyses on any identified clusters from the MVPA analyses. These seed-based analyses were conducted to investigate differences in connectivity patterns between age groups. Individual binary masks of each MVPA cluster were imported as ROIs and used in post-hoc seed-based analyses with between-subject contrast PI High_Old, PI High_Young/Middle-aged (1 −1) and PI Low_Old, PI Low_Young/Middle-aged (1 −1), respectively (voxel *p* = .001 and cluster level *p*^FDR^ = .001).

## 3. RESULTS

### 3.1 Age-related differences and 5-year change in PI

Age was positively correlated with PI scores across age groups (*r*(233) = .339 *p* < .001) at baseline, indicating that increasing age is associated with more PI. Only the younger/middle-aged group showed a significant correlation between age and PI scores at baseline (*r*(127) = .297, *p* = < .001), but not the older group (*r*(102) = .086, *p* = .384). Age was not significantly correlated with change in PI from baseline to follow-up (*r*(130) = .026, *p* =.77), in neither younger/middle-aged (*r*(78) = .022, *p* = .85) or older (*r*(48) = −.133, *p* =.36) adults. PI did not change significantly between baseline and follow-up in the whole sample (*t*(133) = .46, *p* = .65) or within either age-group (younger/middle-aged: *t*(81) = .33, *p* = .74; older: *t*(51) = .32, *p* = .76).

### 3.2 Age-related differences in PI between and within the high and low PI groups

A 2 (age group) × 2 (PI group) ANOVA was performed to evaluate the effects of age group and PI group on PI scores at baseline. The results indicated a significant main effect of age group (*F*(1, 236) =6.38, *p* = .012, partial η^2^ = .03) suggesting higher PI scores in the older group (M = .23, std = .77) than in the younger/middle-aged group (M = −.24, std = .78); a significant main effect for PI group (*F*(1,236) = 395.33, *p* <.001, partial η = .63) suggesting higher PI scores in the high PI group (M = .62, std =.52) than in the low PI group (M =-.67, std = .42); but no interaction between age group and PI group (*F*(1, 236) = .06, p = .81, partial η = .0). The means and standard deviations for baseline PI scores are presented in Table 2.

**Table 2.**
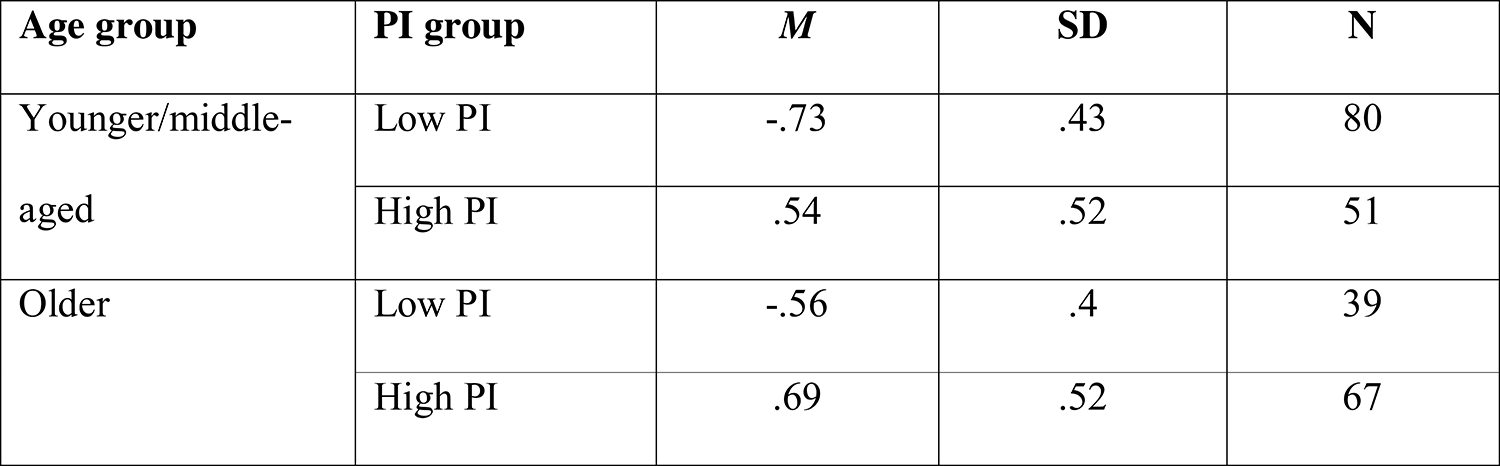
Descriptive statistics for baseline PI scores.

The age effect reached statistical significance in the low PI group (*t*(119) = −2.3, *p* = .023) but not in the high PI group (*t*(117) = −1.58, *p* = .117). The PI effect was significant both in younger/middle-aged adults (*t*(131) = −15.48, *p* <.001) and older adults (*t*(105) = −13, *p* <.001).

### 3.3 Age-related differences in IFG rsFC

Whole brain rsFC analyses with the IFG as a seed region and age group as an independent variable revealed seven clusters showing significant connectivity differences between the age groups (Supplementary table 2). Cluster 1 was centered in the left postcentral gyrus (x, y, z = −36 −20 +40), cluster 2 was centered in the right parahippocampal gyrus (x, y, z = +18 −26 − 16), cluster 3 was centered in the left cuneal cortex (x, y, z = −8 −74 +28), cluster 4 was centered in the right precuneus (x, y, z = +24 −52 +26), cluster 5 was centered in the right superior parietal lobule (x, y, z = +50 −42 +60), cluster 6 was centered in the left supramarginal gyrus (x, y, z = −62 −32 +40), and cluster 7 was centered in the right angular gyrus (x, y, z = +44 −58 +40). Clusters 1, 3, and 7 showed stronger connectivity, whereas clusters 2, 4, 5, and 6 showed weaker connectivity in the older group, compared to the younger/middle-aged group. Analyses of change in IFG connectivity between baseline and follow-up did not reveal any significant differences between the age-groups.

### 3.4 PI is differentially associated with IFG rsFC in younger/middle-aged and older adults

Whole brain rsFC analyses across all participants revealed that higher PI scores were associated with stronger connectivity between the IFG and brain regions within a large cluster centered around the left inferior occipital cortex (x, y, z = **-**32 −80 −10). Full cluster details are reported in Supplementary table 3.

PI was differentially associated with IFG connectivity in younger/middle-aged and older adults in two large clusters (Supplementary table 4 and Figure 1). The first cluster was centered in vermis 4 5( x, y, z = +04 −54 −24) and expanded across parts of the cerebellum and the brain stem. The second cluster was centered in the left caudate (x, y, z = - 04 +14 +10) and expanded across bilateral caudate and the subcallosal cortex. In the vermis cluster, more PI was associated with stronger connectivity in older adults but with weaker connectivity in younger*/*middle-aged adults (Figure 1). In the left caudate cluster, the reverse pattern was observed, with more PI being related to weaker connectivity in older but stronger connectivity in younger*/*middle-aged adults (Figure 1).

**Figure 1.**
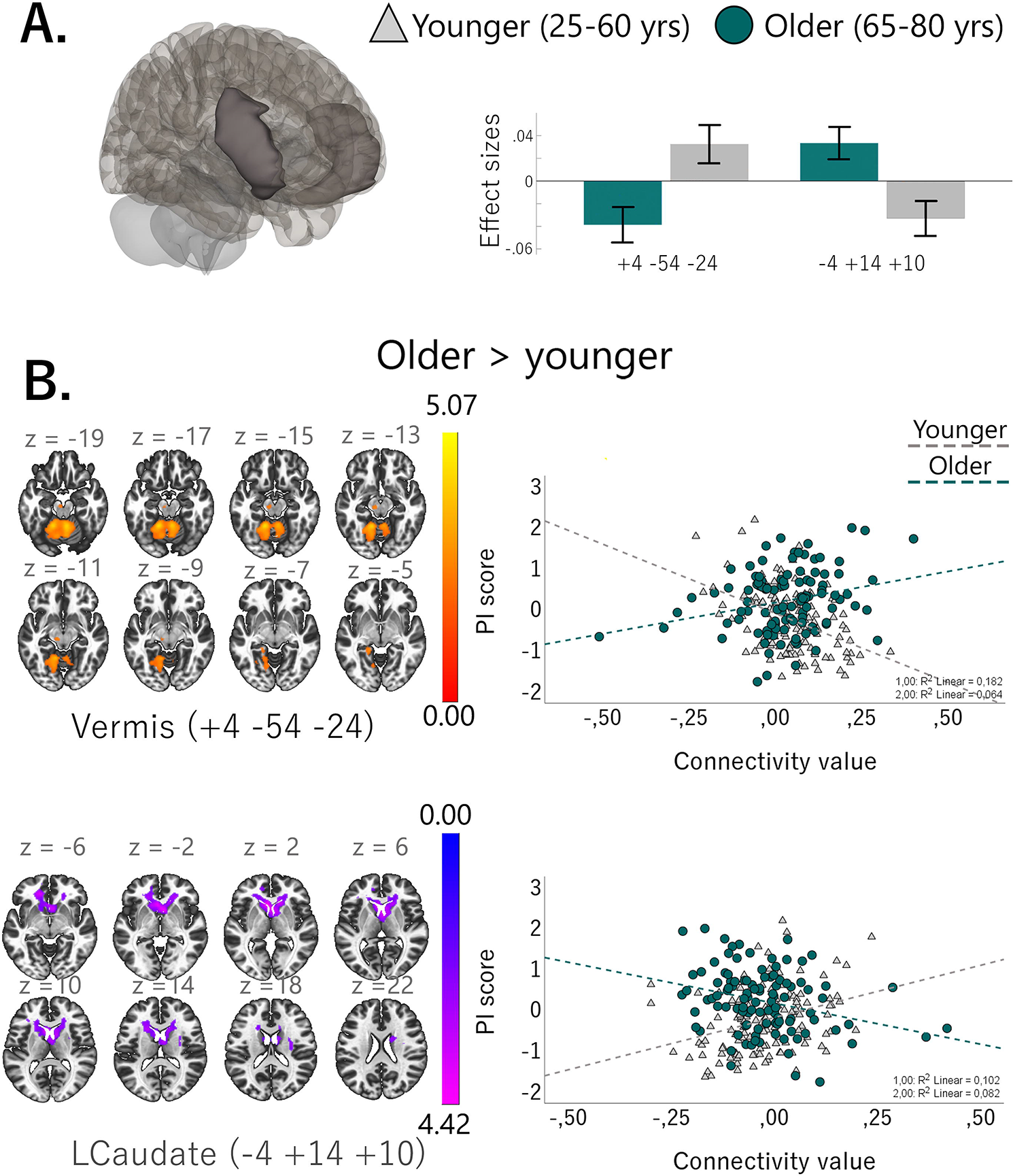
**A**: IFG seed region (to the left) and effect sizes for significant cluster (to the right), **B**: Heat maps for the Older > Younger/middle-aged contrast (to the left) and scatter plots depicting PI score x connectivity value interaction (to the right) in the vermis (top) and left caudate (bottom) clusters. Younger/middle-aged adults are represented in light grey and older adults represented in dark green. *Note: Scatter plots depict the mean connectivity value for each cluster for each participant*.

Analyses of whether 5-year change in PI was associated with 5-year change in IFG rsFC revealed a large (5833 voxels) cluster centered in the left inferior occipital cortex (MNI: −36 −80 −10) spanning several occipital regions (e.g., occipital pole, lingual gyrus, lateral occipital cortex; Supplementary Table 5 and Figure 2). Increased PI was associated with decreased connectivity between the IFG and the inferior occipital cortex cluster across age-groups. Age-group differences in the association between change in PI and change in IFG rsFC were observed in two clusters centered in the right insula (2621 voxels; MNI: +30 +22 +12) and the right superior anterior cingulate cortex (ACC; 2008 voxels; MNI: +12 +30 +24) with older adults showing decreased rsFC between the IFG and these clusters with increased PI and younger/middle-aged adults showing increased rsFC with these clusters with increased PI (Supplementary table 6 and Figure 2).

**Figure 2.**
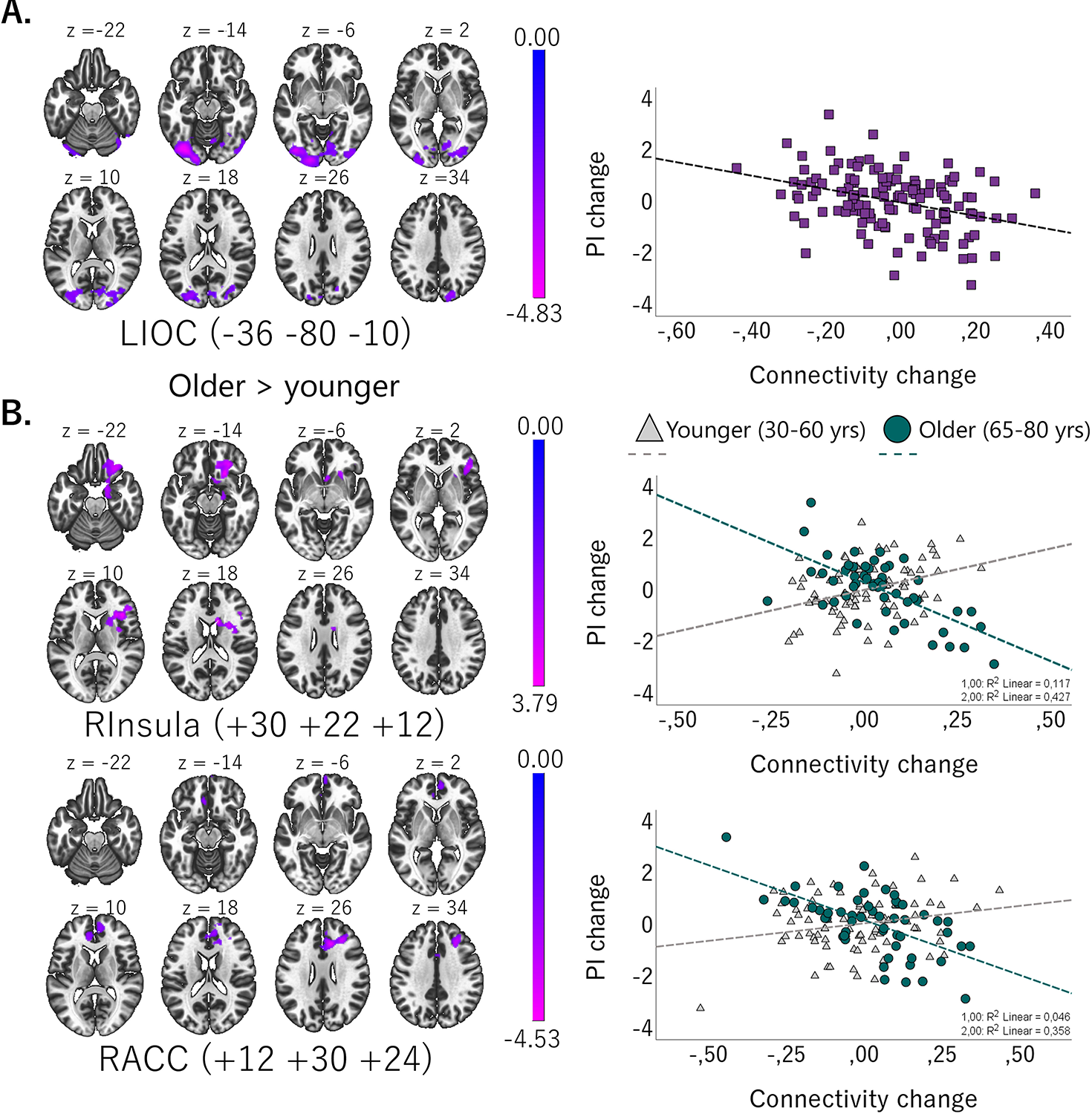
**A**: Results from Change (PI) – Change (IFG rsFC) analyses for the whole sample, including heat maps (left) and scatter plots (right) depicting PI scores x connectivity values. **B:** Results from Change (PI) – Change (IFG rsFC) analyses of age-group differences, including heat maps for the Older > Younger/middle-aged contrast (left) and scatter plots depicting PI score x connectivity value interaction (right). Younger/middle-aged adults are represented in light grey and older adults are represented in dark green. *Note:* Scatter plots depict the mean connectivity value for each cluster for each participant.

### 3.5. Age-related differences in whole-brain rsFC patterns associated with PI

The MVPA analysis examining age-group differences in rsFC patterns within the high PI group revealed 14 significant clusters (*p* = .001, cluster-*p*^FDR^ = .001) where older and younger*/*middle-aged adults displayed significantly different rsFC patterns. These clusters were centered in the right temporal pole, left temporal pole, right posterior cingulate gyrus, right parahippocampal gyrus, insula, cerebellum crus 1, left anterior cingulate gyrus, right temporal pole, left calcarine, right parahippocampal gyrus, left parahippocampal gyrus, right supplementary motor area, left calcarine, and left middle temporal gyrus, respectively. Full clusters details are reported in Table 3. Selected clusters are depicted in Figure 3, as well as age-group connectivity maps for post hoc seed based analyses.

**Figure 3.**
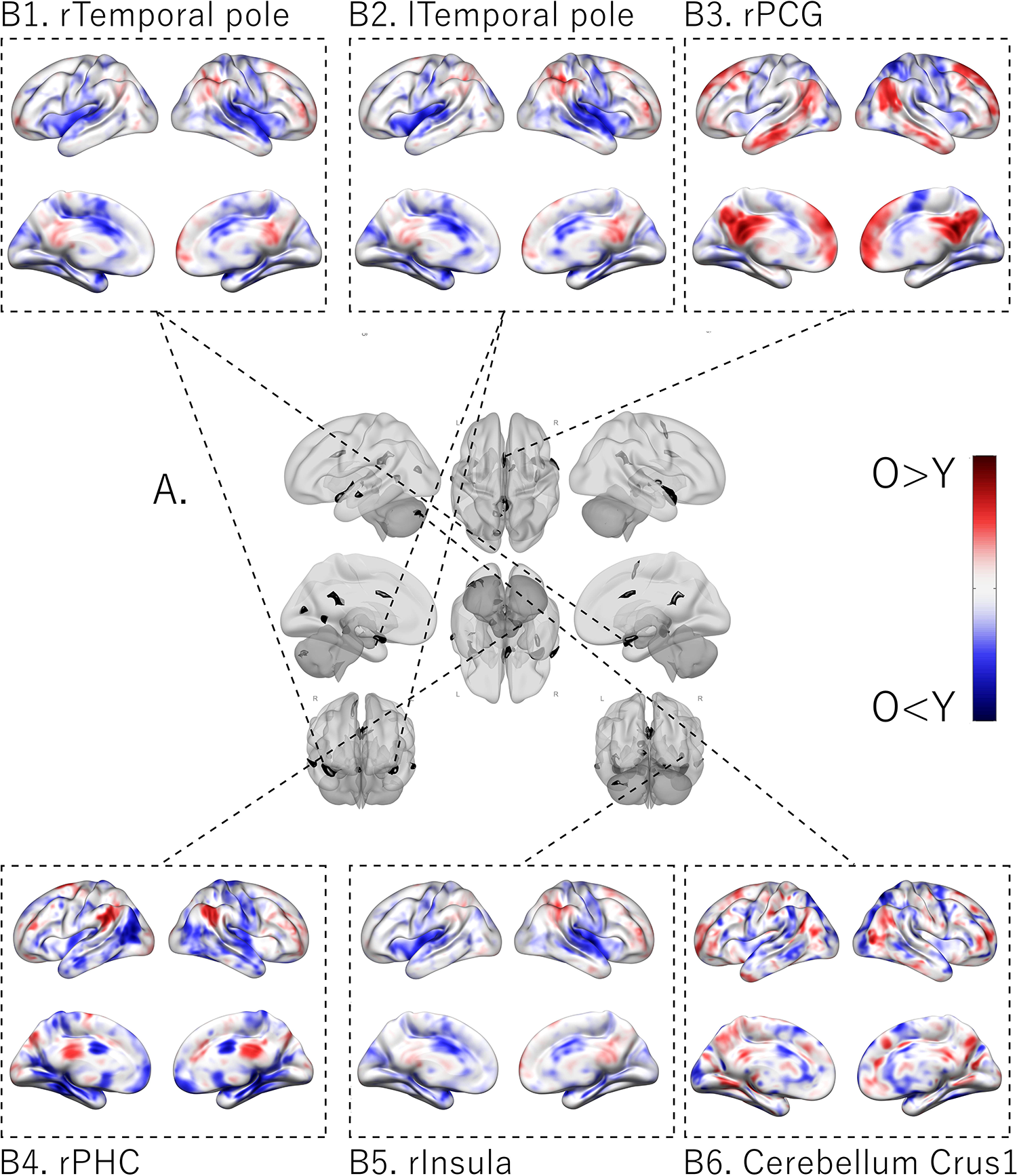
**A.** The central figure (grayscale) shows the clusters (in black) displaying significant age-related differences in functional connectivity in the high PI MVPA (*p*FDR=.001). Depicted here are clusters 1-6 located in the right temporal pole, left temporal pole, right posterior cingulate gyrus, right parahippocampal gyrus, right insula, and cerebellum crus1, respectively. **B.** The patterns of differences between age-groups within each separate cluster, indicated by dashed lines, are depicted at the top (B: 1. Right temporal pole, 2. Left temporal pole, 3. Right posterior cingulate gyrus) and bottom (B: 4. Right parahippocampal gyrus, 5. Right insula, 6. Cerebellum crus1). Red indicates more connectivity in older compared to younger*/*middle-aged participants, blue indicates more connectivity in younger*/*middle-aged compared to older participants. Significant clusters are marked in black. Overall, older adults showed a pattern of weaker connectivity with visual processing regions for the cerebellar, temporal pole, and lingual gyrus seeds. Older adults also displayed weaker connectivity between the cuneus and portions of the temporal, frontal, occipital, and limbic lobes as well as some subcortical regions. Younger*/*middle-aged adults displayed weaker connectivity between the lingual gyrus and prefrontal and occipital regions as well as the superior parietal lobule as well as between the superior temporal gyrus and parietal regions.

The MVPA analysis conducted in the low PI group revealed 6 significant clusters (*p* = .001, cluster-FDR*p* = .001) where older and younger*/*middle-aged adults displayed significantly different rsFC patterns. These clusters were centered in the left postcentral gyrus, right thalamus, right precentral gyrus, left posterior cingulate gyrus, left middle frontal gyrus, and left precentral gyrus, respectively. Full cluster details are reported in Table 4. Clusters are depicted in Figure 4, as well as age-group connectivity maps for post-hoc seed-based analyses.

**Figure 4.**
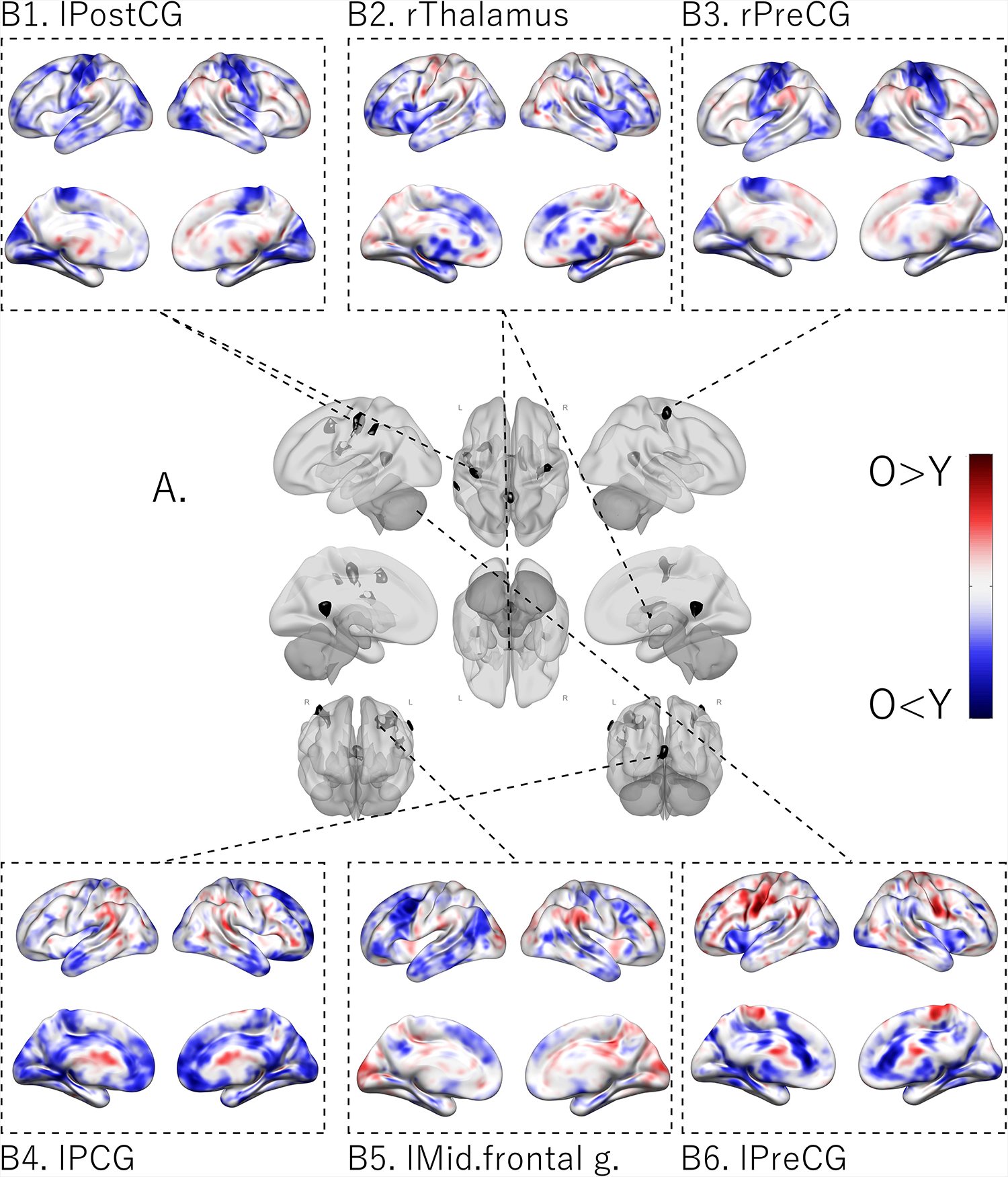
A. The central figure (grayscale) shows the clusters (in black) displaying significant age-related differences in functional connectivity in the low PI MVPA (*p*FDR=.001). These clusters were localized in the left postcentral gyrus, right thalamus, right precentral gyrus, left posterior cingulate gyrus, left middle frontal gyrus, and left precentral gyrus, respectively. **B.** The patterns of differences between age-groups within each separate cluster, indicated by dashed lines, are depicted at the top (B: 1. Left postcentral gyrus, 2. Right thalamus, 3. Right precentral gyrus) and bottom (B: 4. Left posterior cingulate gyrus, 5. Left middle frontal gyrus, 6. Left precentral gyrus). Red indicates more connectivity in older compared to younger*/*middle-aged participants, blue indicates more connectivity in younger*/*middle-aged compared to older participants. Significant clusters are marked in black. Overall, older adults showed stronger connectivity with parietal and frontal regions as well as the insula for both the cerebellum and precentral gyrus seed regions, compared to younger*/*middle-aged adults. Additionally, the precentral gyrus seed showed stronger connectivity with the paracingulate gyrus. Younger/middle-aged adults displayed stronger connectivity with frontal (B2, 3, 4), temporal (B1, 2), limbic (B2, 4), as well as cerebellar and visual processing regions (B2, 4), compared to older adults.

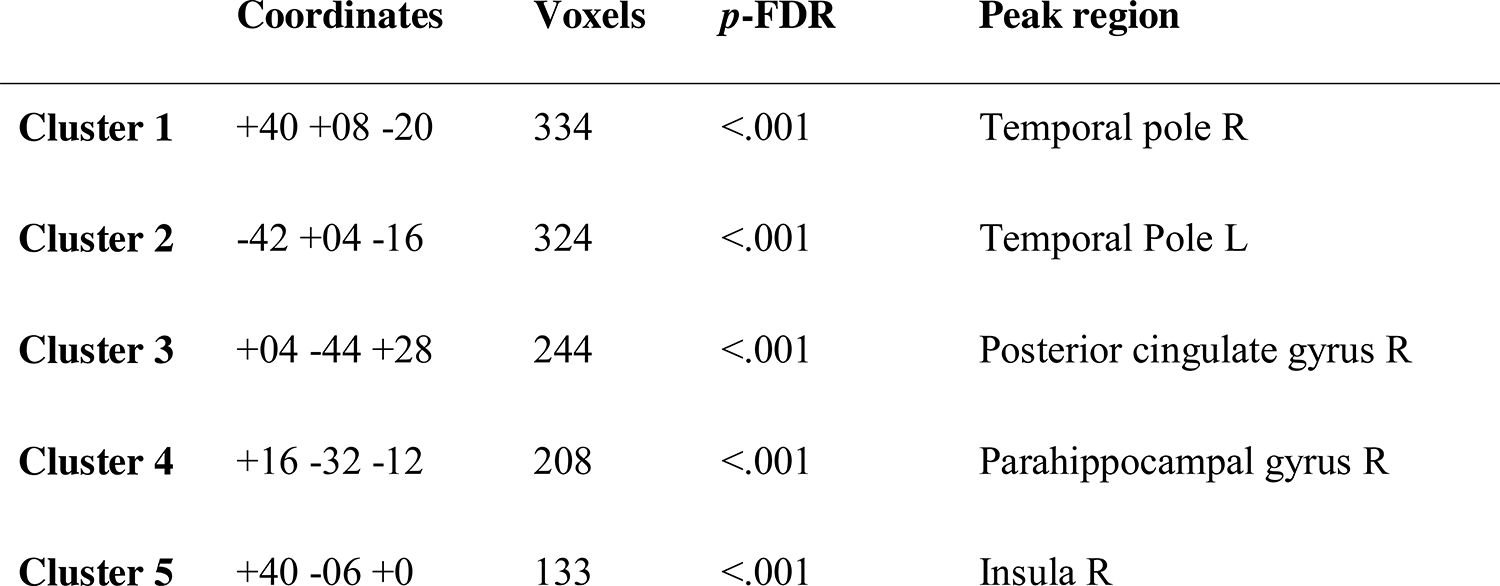
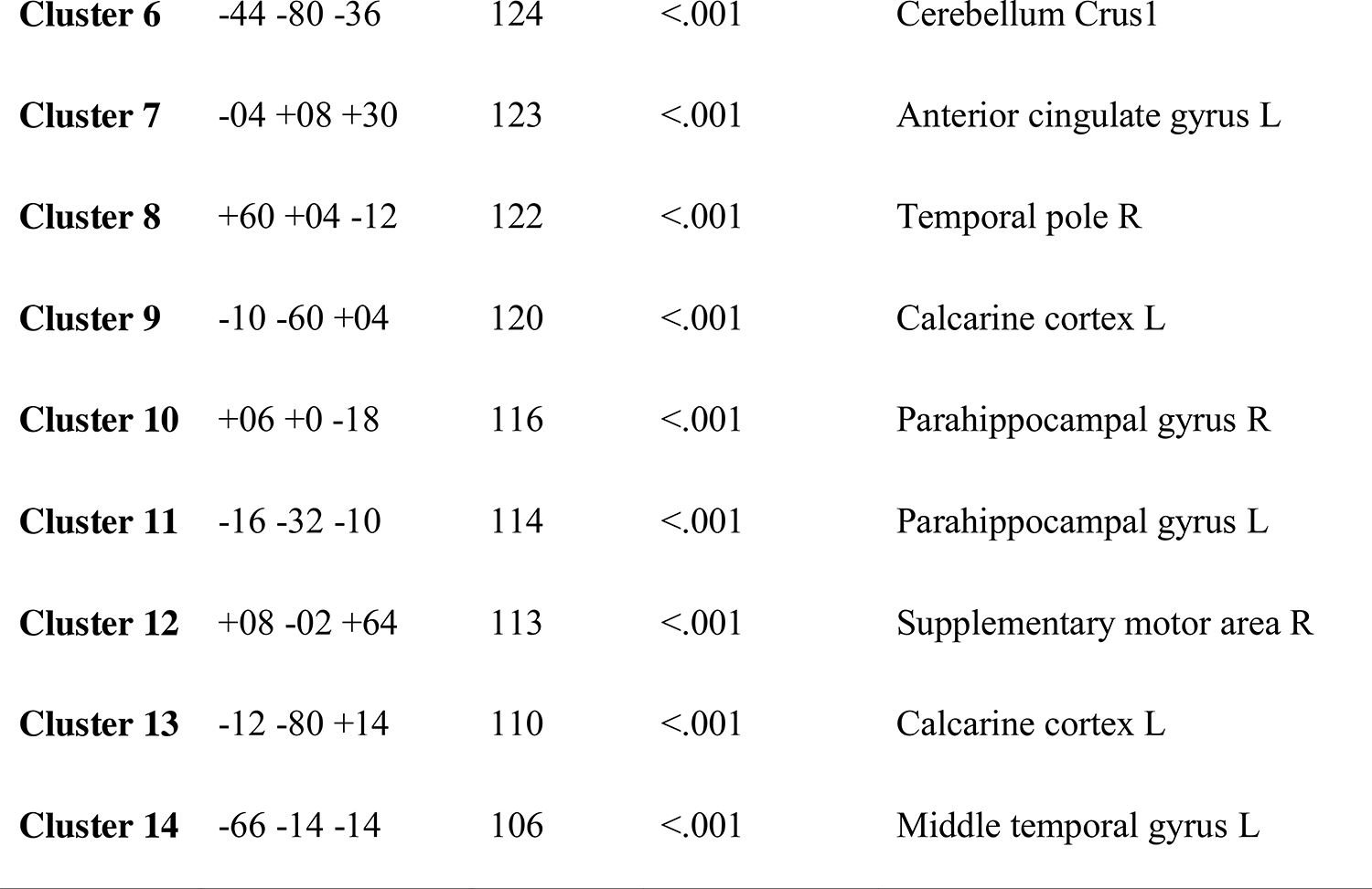

**Table 4.**
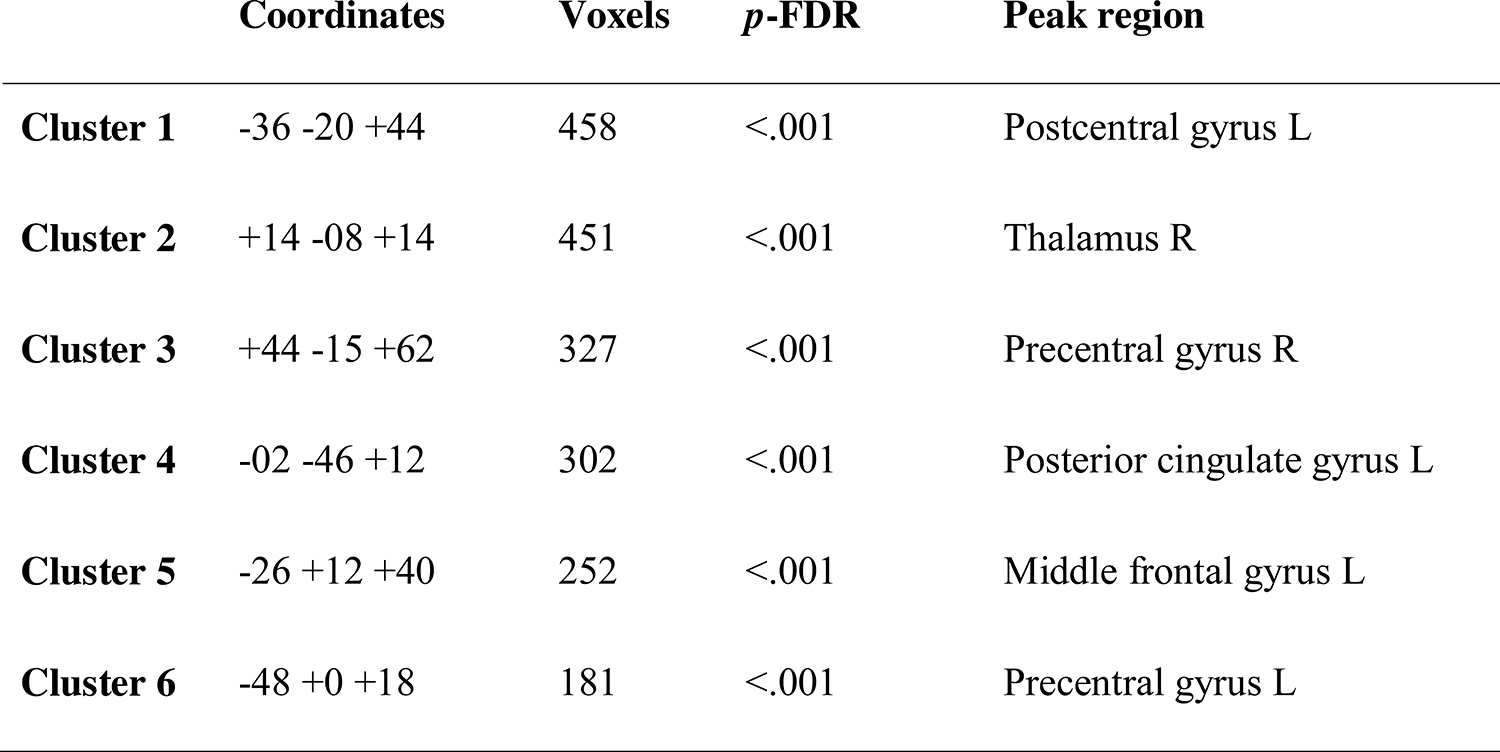
MVPA seeds low PI.

#### 3.5.1 Post-hoc seed-based analyses

Post-hoc seed-based analyses were conducted with the MVPA defined clusters as seed regions. The result of these analyses revealed significant age-group differences in rsFC patterns distributed across cortical and subcortical regions as well as parts of the cerebellum.

In the high PI group, older adults displayed a pattern of weaker connectivity between the right temporal pole with frontal (e.g., postcentral gyrus) and temporal (e.g., Heschl’s gyrus) regions and between the right parahippocampal gyrus with frontal (e.g., postcentral gyrus), temporal (e.g., Heschl’s gyrus), and occipital (cuneal cortex), compared to younger/middle-aged adults. However, one of the right temporal pole seeds additionally showed stronger connectivity with the postcentral gyrus and the hippocampus, compared to younger/middle-aged adults. Similarly, older adults showed weaker connectivity between the left temporal pole with some frontal (e.g., right precentral gyrus) and temporal regions (e.g., right temporal pole), but stronger connectivity with the left precentral gyrus, compared with younger/middle-aged adults. Additionally, older adults showed weaker connectivity between the right insula with temporal (e.g., temporal pole) and limbic (e.g., amygdala) as well as the parahippocampal gyrus but stronger connectivity with the superior frontal gyrus, compared to younger/middle-aged adults.

For the middle temporal gyrus seed, older adults showed weaker connectivity with the left postcentral gyrus and the left inferior temporal gyrus and stronger connectivity with several posterior/visual processing regions (e.g., cerebellum 6, calcarine cortex) compared to younger/middle-aged adults. Older adults also showed stronger connectivity between the posterior cingulate gyrus and the calcarine cortex.

For the right parahippocampal gyrus seed, older adults displayed weaker connectivity with frontal (e.g., IFG triangularis), temporal (e.g., inferior temporal gyrus), and parietal (e.g., angular gyrus) regions, and stronger connectivity with the caudate and parts of the cerebellum, compared to younger/middle-aged adults. For the left parahippocampal gyrus, older adults displayed weaker connectivity with frontal (e.g., insula), parietal (e.g., superior parietal gyrus), temporal (e.g., temporal fusiform gyrus), and parts of the cerebellum coupled with stronger connectivity with other parts of the cerebellum as well as with the caudate and inferior temporal gyrus.

For the anterior cingulate gyrus seed, older adults displayed weaker connectivity with temporal (e.g., superior temporal gyrus), frontal (e.g., IFG triangularis), and cerebellar regions but stronger connectivity with other parts of the IFG and the superior temporal gyrus as well as the caudate, compared to younger/middle-aged adults.

For the left calcarine seeds, older adults showed weaker connectivity with the right superior parietal gyrus and the right supplementary motor area and stronger connectivity spanning frontal (e.g., superior frontal gyrus), temporal (e.g., middle temporal gyrus), parietal (e.g., angular gyrus) regions as well as the cerebellum, compared to younger/middle-aged adults. For the second calcarine seed (−12 −80 +14), older adults displayed stronger connectivity spanning frontal (e.g., IFG orbitalis), temporal (e.g., middle temporal gyrus), and parietal (e.g., angular gyrus) regions.

For the cerebellum seed, older adults displayed weaker connectivity with frontal (e.g., middle frontal gyrus), cerebellum crus2, and the putamen but stronger connectivity with temporal (e.g., Heschl’s gyrus), cerebellum 3, parahippocampal gyrus, angular gyrus, and gyrus rectus, compared to younger/middle-aged adults. Finally, for the supplementary motor area seed, older adults showed weaker connectivity with the calcarine cortex but stronger connectivity with the inferior parietal gyrus, compared to younger/middle-aged adults.

In the low PI group, older adults displayed a pattern of weaker connectivity between the middle frontal gyrus with the bilateral middle temporal gyri and the inferior occipital gyrus, compared to younger/middle-aged adults.

For the postcentral gyrus seed, older adults displayed weaker connectivity with frontal (e.g., postcentral and precentral gyri) regions, the temporal pole, and the inferior occipital gyri but stronger connectivity with other frontal (e.g., insula) regions, compared with younger/middle-aged adults. Similarly, for the left precentral gyrus seed, older adults displayed weaker connectivity with occipital (middle occipital gyrus), temporal (inferior temporal gyrus), medial temporal (parahippocampal gyri), and posterior (vermis 4 5) regions but stronger connectivity with other frontal (e.g., IFG opercularis), compared to younger/middle-aged adults. Moreover, for the thalamus seed, older adults displayed weaker connectivity with frontal (e.g., IFG triangularis) regions as well as with parts of the cerebellum and thalamus and the olfactory but more connectivity with other frontal (e.g., precentral gyri) regions as well as the caudate and the hippocampus, compared to younger/middle-aged adults.

For the right precentral gyrus seed, older adults displayed weaker connectivity with frontal (e.g., precentral gyrus), occipital (e.g., lingual gyrus), and temporal (middle temporal gyrus) but stronger connectivity with the inferior parietal gyrus, compared to younger/middle-aged adults. Older adults also displayed weaker connectivity between the posterior cingulate gyrus seed within the regions as well as with the angular gyrus but stronger connectivity between this seed and the cuneal cortex.

## 4. DISCUSSION

The present study investigated how rsFC is related to PI in WM and how this relationship differed between younger*/*middle-aged and older adults. Both a hypothesis-driven seed-based approach was used to examine the role of IFG rsFC in control of PI, alongside a data-driven MVPA analysis to investigate how the ability to control PI in WM is associated with whole brain connectivity patterns. Of primary interest was to examine whether rsFC – PI associations differed between younger*/*middle-aged and older adults.

The major findings from the study were that 1) older adults showed lower IFG rsFC, cross-sectionally, as compared to younger/middle-aged adults; 2) more PI was associated with stronger IFG rsFC with the left inferior occipital cortex across the whole sample. Additionally, reduced IFG – left inferior occipital cortex connectivity over 5 years was associated with concurrent decline in the ability to control PI; 3) PI showed age-differential relationships in IFG rsFC with the left caudate and the vermis cross-sectionally and with the insula and the anterior cingulate cortex longitudinally; and 4) MVPA analyses identified age-differential patterns of connectivity contributing to the ability to control PI in WM. These novel findings provide converging evidence for altered rsFC with increasing age and confirm that alterations in brain connectivity might have implications for the ability to control interference in WM.

Cross-sectional analyses at baseline across the whole sample revealed stronger connectivity between the IFG and a cluster centered in the left inferior occipital cortex and that weaker connectivity between these two regions was related to more PI. Moreover, longitudinal change–change analyses demonstrated that reduced connectivity between IFG and inferior occipital cortex was associated with reduced ability to control PI across the whole sample.

Furthermore, age stratified analyses revealed that weaker IFG connectivity with the left caudate and stronger IFG connectivity with the vermis were associated with more PI in older adults. The caudate is part of the dorsal striatum of the basal ganglia and has previously been implicated in WM (Lewis et al., 2004; McNab & Klingberg, 2008; Owen et al., 1996; Postle & D’Esposito, 1999). For instance, greater activation of the caudate/striatum has been linked to greater WM capacity and suggested to play a role in controlling attention and filtering distractions in WM (McNab & Klingberg, 2008). Moreover, dopamine depletion in the caudate has been associated with impairments in updating and maintenance of information in WM, indicating the importance of the caudate in dopaminergic pathways involved in cognitive control processes (Cools et al., 2008). Furthermore, caudate dopamine has been associated with IFG activity during WM maintenance (Landau et al., 2009). Computational models have suggested that dopaminergic modulation of activity in the cortico-striatal network mediates executive functioning (e.g., O’Reilly & Frank, 2006). On a related note, Li, Bäckman, and Persson (2019) found that genetic markers associated with dopamine D2 receptor density were associated with frontostriatal activity and WM updating performance, especially in older age.

In a similar vein, previous studies have found that connections between the IFG and the caudate are important in WM updating, and that older adults show reduced coupling between these regions together with reduced performance (Podell, 2012). This functional coupling may reflect inhibitory control processing involved in successfully resolving PI during WM updating. The observed association between more PI and less IFG – caudate rsFC in older adults may, thus, indicate that aberrant IFG – caudate functional connectivity underlies impaired ability to control PI in older individuals.

The possible contribution of the vermis to cognitive functioning, and to WM functioning more specifically, is less clear. Several previous studies have implicated the cerebellum in WM processing (e.g., Cabeza & Nyberg, 2000; Chen & Desmond, 2005; Desmond & Fiez, 1998). However, others have found that while vermeal gray matter volume is associated with cognitive functioning, these associations are no longer significant after controlling for prefrontal volume (Grieve et al., 2009). It has also been suggested that the cerebellum contributes to WM by supporting articulatory rehearsal, postulated to help to maintain information in WM (Ben-Yehudah et al., 2007), which may be important in updating tasks such as the n-back task.

Longitudinal analysis investigating age-group differences in associations between change in IFG rsFC and change in PI across 5 years revealed that a reduced ability to control PI in older adults was associated with reduced connectivity between the IFG and two clusters centered in the insula and the anterior cingulate gyrus, respectively. These regions have previously been implicated in controlling PI in WM (Jonides & Nee, 2006; Loosli et al., 2016; Nee et al., 2007; Samrani & Persson, 2022). The anterior cingulate gyrus has previously been implicated in WM and postulated to play a role of monitoring and resolving conflict, thereby supporting updating and maintenance of relevant information (Botvinick et al., 2004; Smith & Jonides, 1999). Moreover, this region has been implicated in executive functioning more broadly by integrating information from cortical and subcortical regions in order to guide behavior. For instance, Kerns et al. (2004), found that the anterior cingulate cortex is active in tasks requiring conflict monitoring/resolution and this activity was associated with the ability to detect error and adjust behavior. Additionally, Bush et al. (2000) found that the anterior cingulate cortex supports executive functioning through its role in attentional processes and interactions with other frontal regions. Furthermore, older adults have been found to display reduced anterior cingulate activity during WM tasks, indicating reduced efficiency of the region in older age (Madden et al., 2007), and may contribute to age-related declines in WM and executive functioning.

The high PI MVPA analysis showed age-differential patterns of both weaker and stronger functional connectivity. On the one hand, older adults with a lower ability to control PI (that is, low performing older adults) displayed a pattern of weaker connectivity between most of the investigated clusters with frontal and temporal regions of the brain, and also between some seeds with parietal, medial temporal, cerebellar, subcortical, limbic, and occipital regions. Thus, patterns of weaker connectivity were observed globally in older adults compared to younger/middle-aged adults. On the other hand, low performing older adults displayed stronger connectivity between the calcarine cortex, parahippocampal gyrus, temporal pole, and the insula with frontal regions. Notably, patterns of stronger connectivity were not restricted to a specific lobe but were observed more globally, also spanning occipital, cerebellar, parietal, temporal, and medial temporal regions, as well as with the caudate. These age-related differences in connectivity pattern were not attributable to age-differences in PI scores.

The patterns of weaker functional connectivity in older adults may be reflective of age-related structural brain changes, including an anterior-to-posterior gradient of degradation in white matter (Sexton et. al., 2014) and frontal gray matter shrinkage (Fjell et al., 2014). Previous investigations in the Betula sample, based on the same cohort, have found an association between reduced PI and reduced white-matter integrity in aging (Andersson, 2022). Such structural alterations may result in increased activity and/or recruitment of additional regions to compensate for the effect of deterioration on cognitive function (Goh & Park, 2009). For example, weaker rsFC with posterior regions may be compensated by stronger rsFC with frontal regions, consistent with the posterior-anterior shift in aging (PASA; Davis et al., 2008). However, age-differential patterns of weaker connectivity were especially centralized to frontal and temporal regions, and more global patterns were observed across the seed regions. Moreover, patterns of stronger connectivity in low-performing older adults were not centralized to frontal regions and thus, not consistent with a compensatory explanation.

The low PI MVPA analysis also showed age-differential patterns of both weaker and stronger functional connectivity. Older adults with a greater ability to control PI (that is, high performing older adults) displayed a pattern of weaker connectivity between the majority of the investigated seeds with occipital and/or temporal regions. However, weaker connectivity was also observed with some frontal, medial temporal, limbic, and parietal regions, compared to younger/middle-aged adults. In high-performing older adults, compared to younger/middle-aged adults, patterns of stronger connectivity were largely centered in frontal regions, but some seed regions also showed stronger connectivity with parietal and/or occipital regions, as well as the hippocampus and caudate.

Overall, the observed stronger coupling between the left precentral gyrus and the left postcentral gyrus, respectively, with frontal and parietal regions combined with weaker connectivity with posterior/occipital regions, may be reflective of the PASA model (Davis et al., 2008). While both seed regions also displayed patterns of reduced connectivity with some frontal regions, older adults showed stronger connectivity with more frontal regions compared to younger/middle-aged adults. This pattern is consistent with the CRUNCH model (Reuter-Lorenz & Cappell, 2008) and may reflect compensation for other age-related neurocognitive changes (Goh & Park, 2009). While older adults showed significantly more PI than younger/middle-aged adults in the low PI group, they still showed significantly less PI than older adults in the high PI group. Thus, patterns of stronger connectivity in high performing older adults may reflect successful compensation for age-related decline elsewhere. Connectivity patterns consistent with the PASA model were primarily observed within high-performing older adults, consistent with a compensatory mechanism of these alterations. In low-performing older adults, on the other hand, patterns of weaker and stronger connectivity were observed more globally across the brain, with weaker connectivity being more centralized to frontal and temporal regions. Speculatively, this may reflect a greater degree of structural degradation in low performing compared to high performing older adults, an inability to properly engage frontal regions to compensate for age-related decline elsewhere, or some combination of these two. Future studies should address the reasons behind these differences in connectivity by employing a multimodal approach which takes patterns of degradation in gray and white matter into consideration.

While previous studies investigating associations between the ability to control PI and rsFC are scarce, some overlap with the present results and results can be discerned. Samrani & Persson (2022) found that increased distance (5-10 back) between target and lure trials was associated with stronger HC connectivity with the left thalamus and bilateral temporal pole, suggesting the involvement of long-term memory mechanisms in resolving PI. This is consistent with present results from the MVPA analyses demonstrating that low performing older adults display stronger connectivity between the temporal pole and HC and high performing older adults display stronger connectivity between the thalamus and the HC, compared to younger/middle-aged adults. The respective contributions of these functional connections to the ability to control PI in WM is not clear but, speculatively, stronger connectivity between the temporal pole and HC may reflect higher effort for these individuals in retrieving temporal item-context information which rely on the HC. The thalamus has been implicated in WM (e.g., Guo et al., 2017; Inagaki et al., 2019; Samrani et al., 2018) and has been suggested to be more active during low load compared to high load maintenance and has been suggested to contribute to the inhibition of irrelevant information during high load conditions (Gomes et al., 2023). Furthermore, thalamus activity has previously been associated with WM load in the n-back task (Chen, Sorenson, & Hwang, 2023) and that thalamocortical interactions contribute to modulation of distributed cortical activity during WM. Given the important role of the HC in long-term memory, greater connectivity with the HC in older adults may indicate a greater reliance on long-term memory in response to interference conflicts. Furthermore, the HC have been implicated in the ability to control PI in WM and has been suggested to contribute to this ability by providing item-context information about the temporal relevance of information in order to discern goal-relevant from irrelevant information (Beukers et al., 2021; Jonides & Nee, 2006; Oberauer, 2005; Szmalec et al., 2011). The functional implications of differential regional connectivity with the HC in low performing and high performing older adults is not clear but may, speculatively, reflect differences in the utilization of HC-dependent contextual information needed to resolve PI.

Moreover, several brain regions previously implicated in controlling PI (Nee et al., 2007; Samrani & Persson, 2024) were demonstrated to be differentially involved in this ability in younger/middle-aged and older adults in both the IFG-based and whole brain analyses. Seed-based analyses revealed that weaker IFG – caudate connectivity was associated with a lower ability to control PI and that reduced IFG – anterior cingulate gyrus connectivity was associated with reduced ability to control PI in older adults. Moreover, low performing and high performing older adults showed differential connectivity patterns with the caudate. Specifically, low performing older adults showed stronger connectivity between the parahippocampal gyri and the anterior cingulate gyrus with the caudate while high performing older adults showed stronger connectivity between the thalamus with the caudate, compared to younger/middle-aged adults.

### 4.3 Limitations

Important to note is that a relatively liberal voxel-wise and cluster level threshold (.05) was used in the seed-based analyses on IFG rsFC, due to the expected small effect sizes. Consequently, the identified clusters in both the whole sample and age-stratified analyses of the PI effect were large and covered several regions located across different lobes. Thus, interpretations regarding functional implications of these findings should be made with caution as the clusters cover regions also involved in cognitive functions and abilities other than PI in WM.

Longitudinal analyses of IFG rsFC associations with PI revealed a significant cluster centered in the inferior occipital cortex across the whole sample. Moreover, age-differential associations were found between declining ability to control PI and reduced rsFC between the IFG and two clusters centered in the insula and ACC, respectively. Importantly, however, the sample size for the longitudinal analyses were moderate (N = 134). Also, age did not significantly correlate with change in PI scores in the whole sample or within the age-groups. Thus, future studies should aim to replicate the present findings to strengthen conclusions regarding age-related changes in the relationship between PI and IFG rsFC.

### 4.4 Conclusion

We found that older adults displayed stronger IFG rsFC with the postcentral gyrus, cuneal cortex, and angular gyrus but weaker connectivity with the parahippocampal gyrus, precuneus, superior parietal lobule, and supramarginal gyrus at baseline, compared to younger/middle-aged adults.

IFG-based rsFC analyses revealed that weaker IFG connectivity with the left caudate and stronger connectivity with the vermis were associated with more PI in older adults. Moreover, a reduced ability to control PI over 5 years was associated with reduced IFG – inferior occipital cortex connectivity across the whole sample, and reduced IFG – ACC and IFG – insula connectivity in older adults. Additionally, whole brain rsFC analyses revealed age-differential patterns of rsFC associated with a lower and higher ability to control PI in WM. Low performing older adults displayed more global patterns of stronger and weaker connectivity, compared to younger/middle-aged adults, while high performing older adults displayed more centralized patterns of stronger connectivity with frontal regions and weaker connectivity with posterior regions, consistent with models of age-related compensation for neurocognitive decline.

Taken together, these novel results contribute to the understanding of decreased control of PI in aging by showing that both IFG and whole brain rsFC are differentially associated with the ability to control PI in older as compared with younger/middle-aged adults.

## 5. DATA AVAILABILITY

The data that support the findings of this study are available on request from the corresponding author. Due to data protection concerns, publicly sharing the entire data set underlying this study is not possible at the moment.

## 6. AUTHOR CONTRIBUTIONS

Pernilla Andersson: Formal Analysis, Writing - original draft, Writing - review & editing; Martien Schrooten: Supervision, Writing - review & editing and Jonas Persson: Writing - original draft, Writing - review & editing, Supervision, Funding acquisition, Conceptualization.

## 7. FUNDING

This work was supported by the Swedish Research Council (grant number 340-2012-5931), Knut and Alice Wallenberg Foundation (Wallenberg scholar grant to L.N.), Ragnar Söderberg’s Foundation (grant number KVA/2011/88/65 to L.N.), and the Swedish Research Counsil (grant number 2018-01609 to JP.

## 8. DECLARATION OF COMPETING INTERESTS

The authors have no actual or potential conflicts of interest.

## ACKNOWLEDGEMENTS

We acknowledge the contribution by the staff in the Betula project and all participants.

## Notes

### Competing Interest Statement

The authors have declared no competing interest.

