## Supplementary material for "Age differences in brain functional connectivity underlying proactive interference in working memory"

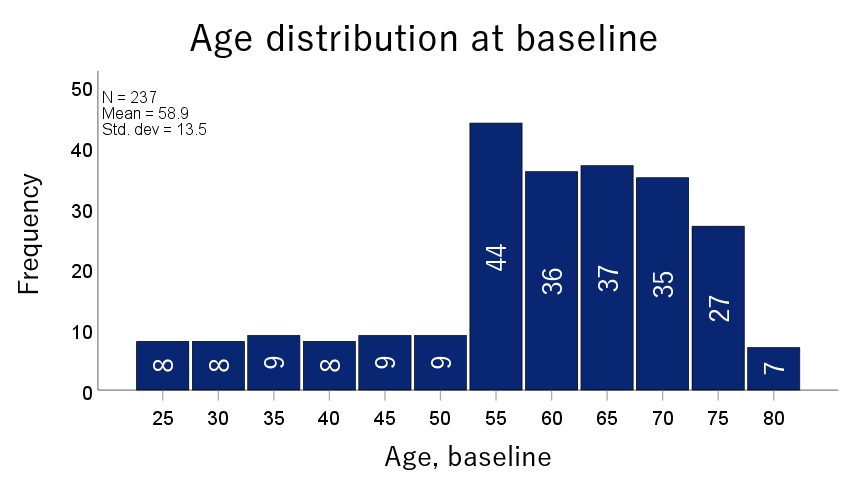

**Supplementary figure 1.** Bar chart displaying the distribution of participants across different age bins at baseline*.*

|  | Baseline | | Follow-up | |
| --- | --- | --- | --- | --- |
|  | LowPI | HighPI | LowPI | HighPI |
| N | 119 | 118 | 69 | 65 |
| Age-group (Y/O) | 80/39 | 51/67 | 49/20 | 33/32 |
| Sex (F/M) | 64/55 | 57/61 | 31/38 | 32/33 |
| Education, in years | 14.1 (4.1) | 13 (3.5) | 14.4 (4.1) | 13.7 (3.4) |
| MMSE | 28.5 (1.3) | 28.3 (1.3) | 28.5 (1.2) | 28.2 (1.5) |
| Combined PI score T5 | -.67 (.42) | .62 (.52) | -.72 (.45) | .69 (.59) |

**Supplementary table 1.** Demographics for the High and Low PI groups.
*Note: Mean (SD); Y/O = Young/Old; F/M = Female/Male; MMSE = Mini mental-state examination; PI = Proactive interference.*

|  | **Coordinates** | **Voxels** | ***p*-FDR** | **Center region** |
| --- | --- | --- | --- | --- |
| **Cluster 1** | **-36 -20 +40** | **5446** | **<.001** | **Postcentral gyrus L** |
| Frontal pole L (28%) | -26 +54 +6 | 1506 |  |  |
| Frontal pole R (14%) | +30 +56 -6 | 762 |  |  |
| Middle frontal gyrus L (13%) | -36 +16 +46 | 693 |  |  |
| Precentral gyrus L (12%) | -44 -1+ +42 | 630 |  |  |
| Paracingulate gyrus L (6%) | -6 +26 +36 | 350 |  |  |
| **Cluster 2** | **+18 -26 -16** | **5043** | **<.001** | **Parahippocampal gyrus R** |
| Brain stem (7%) | +2 -36 -28 | 374 |  |  |
| Temporal pole R (7%) | +46 +10 -18 | 343 |  |  |
| **Cluster 3** | **-8 -74 +28** | **2978** | **.002** | **Cuneal cortex L** |
| Precuneous (35%) | -2 -64 +28 | 1057 |  |  |
| Posterior cingulate gyrus (22%) | +0 -40 +26 | 662 |  |  |
| Cuneal cortex L (10%) | -8 -80 +24 | 283 |  |  |
| Cuneal cortex R (8%) | +8 -78 +26 | 241 |  |  |
| Intracalcarine cortex R (6%) | -6 -78 +10 | 183 |  |  |
| **Cluster 4** | **+24 -52 +26** | **2294** | **.006** | **Precuneus R** |
| Anterior cingulate gyrus (7%) | +0 -2 +30 | 159 |  |  |
| **Cluster 5** | **+50 -42 +60** | **1960** | **.01** | **Superior parietal gyrus R** |
| Supramarginal gyrus, anterior, R (33%) | +60 -28 +38 | 649 |  |  |
| Supramarginal gyrus, posterior R (17%) | +60 -38 +42 | 334 |  |  |
| Postcentral gyrus R (15%) | +62 -18 +36 | 290 |  |  |
| Superior parietal lobule R (8%) | +40 -46 +58 | 151 |  |  |
| **Cluster 6** | **-62 -32 +40** | **1762** | **.02** | **Supramarginal gyrus L** |
| Supramarginal gyrus, anterior L (39%) | -60 -32 +36 | 692 |  |  |
| Postcentral gyrus L (15%) | -60 -22 +32 | 268 |  |  |
| Supramarginal gyrus, posterior L (11%) | -58 -44 +42 | 200 |  |  |
| Superior temporal gyrus L (5%) | -64 -36 +10 | 92 |  |  |
| Parietal operculum L (5%) | -60 -34 +22 | 81 |  |  |
| **Cluster 7** | **+44 -58 +40** | **1416** | **.04** | **Angular gyrus R** |
| Lateral occipital cortex R (57%) | +46 -66 +40 | 809 |  |  |
| Angular gyrus R (36%) | +50 -54 +36 | 515 |  |  |

**Supplementary table 2.** Table depicting resulting clusters identified in IFG rsFC analyses on differences between older and younger/middle-aged adults (older > younger), including (from left to right) coordinates, cluster size, *p*-FDR, and center region. Peak cluster details are bolded. Below, subpeaks are reported, including their respective contribution (in percentage) to the cluster size. *Note: Subpeaks are only reported for regions contributing 5 or more % to total cluster volume.*

| **Cluster region** | | **Voxels** | **Coordinates** | ***p*FDR** |
| --- | --- | --- | --- | --- |
| **Inferior Occipital L** | | **4169** | **-32 -80 -10** | **< .001** |
|  | Lateral Occipital cortex inferior L (15%) | 623 | -32 -78 +24 |  |
|  | Occipital pole L (13%) | 546 | -18 -94 +4 |  |
|  | Cuneal cortex L (8%) | 329 | -8 -80 +26 |  |
|  | Occipital fusiform gyrus L (8%) | 322 | -28 -76 -12 |  |
|  | Intracalcarine L (7%) | 297 | -12 -70 +10 |  |
|  | Lingual gyrus L (6%) | 260 | -14 -66 -4 |  |
|  | Lateral occipital cortex inferior L (6%) | 256 | -36 -84 -4 |  |

**Supplementary table 3.** Table depicting details about the cluster identified from the analysis of PI effect on IFG rsFC at baseline, including (from left) cluster peak region, cluster size, MNI coordinates, and *p*FDR. Cluster details are bolded. Below, details from subpeaks are reported. *Note: Only subpeaks contributing 5% or more to total cluster volume are reported;* *subpeak-region(contribution to cluster volume)*

| **Cluster** | | **Coordinates** | **Size** | ***p*FDR** |
| --- | --- | --- | --- | --- |
| **Vermis 4 5** | | **+04 -54 -24** | **6934** | **<.001** |
|  | Cerebellum 8 R (10%) | +26 -60 -48 | 713 |  |
|  | Cerebellum 6 L (10%) | -18 -60 -22 | 693 |  |
|  | Cerebellum 6 R (8%) | +20 -62 -24 | 572 |  |
|  | Cerebellum crus1 L (8%) | -26 -72 -30 | 556 |  |
|  | Brain stem (5%) | +2 -30 -28 | 381 |  |
|  | Cerebellum 4 5 L (5%) | -12 -50 -18 | 367 |  |
|  | Cerebellum 8 L (5%) | -22 -60 -48 | 334 |  |
| **Left Caudate** | | **-04 +14 +10** | **3539** | **<.001** |
|  | Caudate R (10%) | +12 +12 +8 | 364 |  |
|  | Caudate L (8%) | -12 +14 +8 | 281 |  |
|  | Subcallosal cortex (7%) | +0 +18 -8 | 252 |  |

**Supplementary table 4.** Table depicting details about the clusters from the analyses on Age group x PI interaction effect on IFG rsFC at baseline, including (from left to right) cluster center region, peak coordinates, cluster size, *p*-FDR. Below, details from subpeaks are reported. Region labels correspond to the AAL atlas. *Note: Only subpeaks contributing 5% or more to total cluster volume are reported;* *subpeak-region(contribution to cluster volume)*

| **Cluster region** | | **Voxels** | **Coordinates** | ***p*FDR** |
| --- | --- | --- | --- | --- |
| **Inferior Occipital L** | | **5833** | **-32 -80 -10** | **< .001** |
|  | Occipital pole L (13%) | 739 | -18 -96 -4 |  |
|  | Lateral occipital cortex inferior R (12%) | 676 | +42 -80 -6 |  |
|  | Occipital fusiform gyrus L (10%) | 564 | -28 -80 -14 |  |
|  | Lateral occipital cortex inferior L (8%) | 489 | -38 -82 -10 |  |
|  | Lingual gyrus R (6%) | 321 | +10 -74 -6 |  |
|  | Lateral occipital cortex superior L (5%) | 298 | -28 -84 +18 |  |

**Supplementary table 5.** Table depicting details about the cluster identified from the Change (PI) x Change (IFG rsFC) analyses in the full sample, including (from left) cluster peak region, cluster size, MNI coordinates, and *p*FDR. Cluster details are bolded. Below, details from subpeaks are reported. *Note: Only subpeaks contributing 5% or more to total cluster volume are reported;* *subpeak-region(contribution to cluster volume)*

| **Cluster region** | | **Voxels** | **Coordinates** | ***p*FDR** |
| --- | --- | --- | --- | --- |
| **Insula R** | | **2621** | **+30 +22 +12** | **.007** |
|  | Orbitofrontal cortex R (18%) | 468 | +24 +24 -18 |  |
|  | Frontal pole R (9%) | 246 | +30 +38 -12 |  |
|  | Frontal operculum R (6%) | 147 | +38 +22 +6 |  |
| **Anterior cingulate gyrus R** | | **2008** | **+12 +30 +24** | **.02** |
|  | Frontal pole R (21%) | 423 | +18 +56 +18 |  |
|  | Anterior cingulate gyrus (20%) | 410 | +4 +32 +20 |  |
|  | Paracingulate gyrus R (18%) | 366 | +8 +46 +14 |  |
|  | Middle frontal gyrus R (14%) | 286 | +30 +30 +40 |  |

**Supplementary table 6.** Table depicting details about the clusters identified from the age-group contrasted Change (PI) x Change (IFG rsFC) analyses, including (from left) cluster peak region, cluster size, MNI coordinates, and *p*FDR. Cluster details are bolded. Below, details from subpeaks are reported. *Note: Only subpeaks contributing 5% or more to total cluster volume are reported;* *subpeak-region(contribution to cluster volume)*
